# Spatial transcriptomics maps region and isoform specific brain alterations in *mdx* and *mdx^4cv^* models of Duchenne muscular dystrophy

**DOI:** 10.64898/2026.10.13.758478

**Authors:** Qirong Mao, Laura GM Heezen, Sharon de Vries, Nynke Verwoerd, Annemieke Aartsma-Rus, Maaike van Putten, Pietro Spitali, Ahmed Mahfouz

## Abstract

Duchenne muscular dystrophy (DMD) is accompanied by cognitive and neurodevelopmental difficulties. To investigate how the loss of different dystrophin isoforms affects gene expression across brain regions, we performed spatial transcriptomics on brains from *mdx* and *mdx^4cv^* mouse models (lacking Dp427 or Dp427+Dp140 respectively) and healthy controls aged 2 weeks. Both dystrophic models showed transcriptional changes converging on myelin and oligodendrocyte biology and oxidative phosphorylation, but with distinct regional patterns. In *mdx* mice, myelin genes were upregulated in the amygdala and hippocampus and downregulated in fiber tract regions, while mitochondrial genes were upregulated in most regions but downregulated in the thalamus and isocortex. In *mdx^4cv^* mice, myelin genes were broadly upregulated and mitochondrial genes downregulated across almost all regions. Direct comparison between *mdx* and *mdx^4cv^* mice demonstrated that the additional loss of Dp140 expanded localized myelin upregulation across the brain and shifts the mitochondrial response toward widespread downregulation. These differences in regional changes between the mouse models is in line with their dystrophin isoform(s) loss. Our findings provide a molecular basis for the brain-related impairments of DMD and a foundation for future therapeutic studies.

## Introduction

Duchenne muscular dystrophy (DMD) is an X-linked recessive neuromuscular disorder caused by out-of-frame mutations in the *DMD* gene, resulting in the absence of functional dystrophin. It is one of the most common and severe forms of muscular dystrophy, affecting approximately 1 in 5,000 male newborns (1). The disease is characterized by progressive muscle wasting and weakness, leading to loss of ambulation and reduced life expectancy. Beyond muscle pathology, brain-related impairments have also been reported in individuals with DMD. The mean Full-Scale Intelligence Quotient (FSIQ) in individuals with DMD is about one standard deviation below that of the general population, and about one-third of individuals with DMD have cognitive impairment in the intellectual disability range (FSIQ < 70) (2-4). Among individuals with DMD, an increased prevalence of attention-deficit/hyperactivity disorder (ADHD), autism spectrum disorder (ASD), obsessive-compulsive disorder (OCD), emotional disorders, and academic difficulties has been reported (5-10). As improvements in standards of care have extended life expectancy, cognitive and behavioral impairments require increasing attention to preserve quality of life (11, 12). Unlike its role in maintaining membrane stability in muscle, dystrophin’s functions in the brain are more complex and isoform-specific during neurodevelopment. No approved therapies currently target the underlying molecular mechanisms of DMD associated cognitive and neurodevelopmental disorders largely because these processes remain poorly defined, and current management is mainly symptomatic. Clinical reviews indicate that psychopharmacological interventions, particularly methylphenidate, offer meaningful clinical improvement in managing comorbid ADHD in patients with Duchenne muscular dystrophy (13).

The *DMD* gene is one of the largest genes in the human genome and is notable for its genomic complexity, utilizing at least seven promoters to generate a diverse repertoire of isoforms expressed in the brain. These include the full-length isoforms Dp427c and Dp427p, as well as shorter variants such as Dp140, Dp71, and Dp40, which show different spatial-temporal expression patterns during neurodevelopment (14, 15). However, dissecting the individual contributions of these isoforms across multiple brain regions has been limited by transcriptomic approaches that lack the spatial resolution needed to distinguish region-specific effects.

Moreover, most transcriptomic studies utilize the classic *mdx* mouse, which lacks the full-length Dp427 isoforms but retains the shorter Dp140, Dp71, and Dp40 isoforms (16). Consequently, it fails to recapitulate the more severe cognitive phenotypes associated with distal mutations additionally disrupting shorter isoforms (17).

To investigate the consequences of dystrophin isoform loss for gene expression across brain regions, we studied transcriptomic profiles in mice with different mutation types, including *mdx* mice lacking the full-length Dp427 isoforms, and *mdx^4cv^* mice lacking both Dp427 and Dp140 isoforms. We generated spatial gene expression profiles of the DMD mouse brain using spatial transcriptomics (10x Genomics Visium). Using this data, we investigated differences in gene expression associated with Dp427 and Dp140 deficiency across brain regions. Our findings reveal distinct regional changes in myelin-related and oxidative phosphorylation gene expression depending on dystrophin isoform deficiency. In *mdx* mice, myelin-associated genes were upregulated in the hippocampus and amygdala but downregulated in fiber tracts, whereas oxidative phosphorylation genes were upregulated in most regions except the thalamus and isocortex. In *mdx^4cv^* mice, the additional loss of Dp140 was associated with widespread myelin-related gene upregulation and oxidative phosphorylation gene downregulation, suggesting broader transcriptional alterations across the brain. Together, these findings help bridge knowledge gaps in our understanding of the molecular mechanisms underlying brain-related impairments in DMD.

## Methods and Materials

### Animals

This study included male C57BL/6J Wildtype (WT; *n* = 3 mice), *mdx* (*n* = 3 mice), and *mdx^4cv^* (*n* = 3 mice) mice, all on a C57BL/6J genetic background (18, 19). Four WT tissue sections were analyzed because one WT mouse contributed two sections. Mice were bred at the animal facility of the Leiden University Medical Center (LUMC). The experiments were approved by the Animal Ethics Committee of the LUMC (AVD 1160020171407, PE.17.246.004) and executed in accordance with Directive 2010/63/EU of the European Parliament.

### Visium sample preparation

For brain sample collection, mice (postnatal day 14 - 17) were deeply anesthetized with an overdose of isoflurane and immediately transcardially perfused with ice-cold artificial cerebrospinal fluid (aCSF: 87 mM NaCl, 2.5 mM KCl, 1.25 mM NaH₂PO₄, 26 mM NaHCO₃, 75 mM sucrose, 20 mM glucose, 2 mM CaCl₂, and 2 mM MgSO₄) following a previously reported procedure (20). Mice were decapitated, and brains were rapidly dissected into ice-cold aCSF. The brains were subsequently rapidly frozen in chilled isopentane and stored at −80°C until further processing.

Following a 30 min equilibration period at cryostat chamber temperature (−20°C), brains were embedded in Tissue-Tek O.C.T. Compound (Sakura Finetek, Torrance, CA, USA) and rapidly frozen using the fast-freezing function of the cryostat object cooler. Tissues were cryosectioned at 10 μm using the Leica CM3050 S cryostat (Leica Microsystems, Wetzlar, Germany) in sagittal orientation and sections were placed on Visium Spatial Gene Expression slides (PN: 2000233, 10X Genomics). Sections were adhered by briefly warming the back of the slide and each capture area only included a single section. Slides were stored at -80°C until continuation with methanol fixation and H&E staining according to the 10X Genomics protocol (CG000160) Images of the capture areas were acquired using a ZEISS Axio Scan.Z1 slide scanner (Carl Zeiss Ltd, Cambridge, UK) with a 20× objective, and exported as TIFF files for alignment to the sequencing data. After imaging spatial gene expression libraries were generated by subsequently permeabilizing the tissue to capture mRNA onto the Visium slide, generating full-length cDNA through reverse transcriptase with template-switching oligonucleotides, second strand synthesizing, denaturizing and amplifying the cDNA, fragmentizing and size-selecting cDNA and indexing according to the manufacturer’s instructions (10X Genomics, CG000239). cDNA amplification used 12-17 cycles, determined from Cq values. Spatial gene expression libraries were sequenced on an Illumina NovaSeq 6000 (Illumina Inc., San Diego, CA, USA) with sequencing depth determined according to the number of tissue-covered spots and targeting approximately 50,000 paired-end reads per covered spot.

### Visium data preprocessing and analysis

Visium spatial transcriptomics data were analyzed using Scanpy v1.9.1 and scvi-tools v1.1.4 in Python v3.10 (21, 22). For the first batch of data, ribosomal (*Rps*, *Rpl*), mitochondrial (*Mt*-), and hemoglobin (*Hba-*, *Hbb-*) geneswere removed from the expression matrix, and cerebellar spots were manually excluded based on their anatomical location and consistently low transcript counts observed in the Space Ranger v2.0.1. The remaining spots from the first Visium batch were then used to generate an unsupervised latent representation was generated using scVI implemented in scvi-tools with default model parameters. A k-nearest-neighbor graph was constructed from the scVI latent space using k = 15 neighbors, followed by Leiden clustering at a resolution of 1.0. The resulting clusters were annotated as spatial brain regions based on established anatomical structures, visual inspection of their spatial distribution, and region-specific marker genes derived from the Allen Mouse Brain Atlas. The annotated spots from the first batch were subsequently used as labelled reference data for label transfer to the unannotated second-batch Visium data using scANVI implemented in scvi-tools. The scANVI model was initialized from the pretrained scVI model, using annotation as the label key and Unpredicted as the unlabeled category. The model was trained using 70% of the data for training and 30% for validation, with other training parameters set to the scvi-tools defaults. Predicted scANVI labels were subsequently used to assign spatial-domain annotations to the second-batch spots.

Data preprocessing and quality control (QC) were performed using the Seurat v5.0.3 in R v4.3.2 following the initial annotation (23, 24). For each spatial spot, QC metrics were calculated, including total unique molecular identifier (UMI) counts, total detected features, and the proportional transcript abundance of mitochondrial and hemoglobin genes. To remove technical artifacts, multiplets, and low-quality spots, global upper-limit thresholds were applied across all spatial regions: spots exceeding 35,000 UMI counts, 6,500 detected features, or a 5% hemoglobin transcript proportion were excluded from downstream analysis. Furthermore, to account for anatomical differences in baseline, cellular density and transcript yield, region-specific viability thresholds were implemented based on manual annotations. Spots mapping to the hippocampus formation were retained if they exhibited > 200 UMI counts, > 100 detected features, and < 40% mitochondrial transcripts. Spots located outside the hippocampus were subjected to more stringent criteria, requiring > 500 UMI counts, > 300 detected features, and < 25% mitochondrial transcripts. We also removed the whole cerebellum region due to the low number of reads in the region. Following spot-level filtering, highly abundant structural and metabolic genes such as specifically ribosomal (*Rps*, *Rpl*), mitochondrial (*Mt*-), and hemoglobin (*Hba-*, *Hbb-*) genes, were computationally removed from the primary spatial assay to prevent them from disproportionately influencing downstream variance, dimensional reduction, and clustering.

### Differential expression analysis

Differential gene expression analysis was performed separately within each annotated brain region using MAST v1.28.0 in R (25). For each brain region, pairwise comparisons were conducted between mice genotype groups using log-normalized gene expression values. MAST applies a two-part hurdle model that separately models the probability of gene detection and the continuous expression level conditional on detection, making it suitable for sparse spatial transcriptomic data. For each gene, expression was modeled as a function of the mice genotype group while adjusting for experimental batch and the scaled number of detected genes per spot *cngeneson* as covariates:

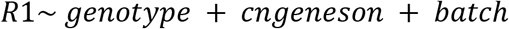

The latter was included to account for differences in gene detection across spots, while batch was included to adjust for technical variation between experimental batches. Only genes detected in more than 10% of spots within each comparison were included in the analysis. Differential expression was assessed using likelihood-ratio tests, and multiple-testing correction was performed using the Benjamini–Hochberg method.

### Enrichment analysis

GO enrichment analyses were performed using the clusterProfiler v4.10.1 in R (26). Differentially expressed genes were mapped to Entrez gene identifiers using the mouse annotation database. Over-representation analysis was conducted with enrichGO() for the Biological Process ontologies, using all genes in Visium samples passing expression filtering as the background universe. *P* values were adjusted using the Benjamini–Hochberg method, and GO terms with adjusted *P* < 0.05 were considered significant. Redundant GO terms were removed using the simplify function with a semantic similarity cutoff of 0.7, retaining the terms with the lowest adjusted *P* value among similar terms. GSEA was performed using gseGO() on all tested genes ranked by the covariate from MAST.

### Neurodevelopmental disorder related genes in SysNDD database

Differential expression results were further evaluated for overlap with neurodevelopmental disorder associated genes using the SysNDD database (27). SysNDD is a curated resource of genes implicated in neurodevelopmental disorders and assigns genes to evidence-based categories according to the strength of genetic associations. We used these annotations to define a curated set of genes that SysNDD categorizes as having ‘established’, ‘strong’, or ‘moderate’ evidence for involvement in neurodevelopmental phenotypes.

### Inference of Transcription Factor Activity and Cross-Strain Comparison

To infer changes in transcription factor (TF) activity across different brain regions and genetic models, we utilized the decoupleR v2.9.7 in conjunction with the comprehensive CollecTRI gene regulatory network for mice (28). Prior to analysis, differential gene expression statistics derived from MAST were transformed into a continuous transcriptomic signature. For each gene, a signed significance metric was calculated by multiplying the sign of the log-fold change by the negative log10-transformed False Discovery Rate (FDR). Scores resulting in positive or negative infinity due to FDR values of zero were capped at +300 and −300, respectively. Annotated regions with very low spots (hindbrain, meninges, and ventricles) were excluded from this spatial analysis. We then estimated TF activities for each contrast using the Univariate Linear Model algorithm (run_ulm), restricting the inference to TFs with a minimum of 5 annotated downstream target genes. To determine the extent of disease-driven transcriptional (dys)regulation shared between the two dystrophic mouse models, we performed a cross-strain exclusivity analysis within each spatial region. Inferred TFs were classified into distinct functional categories based on their ULM score directionality and statistical significance (*P* < 0.05).

### Module score calculation of mitochondrial electron transport chain complex

To spatially resolve the activity of the ETC, we curated complex-specific mitochondrial gene sets utilizing the MitoCarta3.0 database (29). Specifically, genes encoding subunits and associated functional factors of the distinct ETC complexes (Complex I–V) were extracted and mapped to our filtered dataset. To quantify the collective expression of these overlapping ETC components at the individual spot level, we computed composite gene signature scores for each complex using the AddModuleScore() function in Seurat. This approach allowed us to evaluate the relative enrichment and spatial distribution of specific ETC metabolic programs across the tissue architecture.

## Results

### Spatial transcriptomics mapping of gene expression in the DMD mouse brain

To investigate spatial transcriptomic variation associated with the loss of full-length Dp427 isoforms and the shorter Dp140 isoform, we performed Visium spatial transcriptomics on sagittal brain sections from wildtype (WT), *mdx*, and *mdx^4cv^* mice at approximately postnatal day 14 (Figure 1a). First, we performed unsupervised clustering of Visium spots from slide 1, which was generated in the first experimental batch, and annotated the resulting clusters by matching anatomical information and region-specific marker-gene expression to the Allen Brain Atlas (30). We then integrated slides 2 and 3, which were generated in a second experimental batch, with slide 1 and transferred the brain-region labels to these slides (Figure 1b). After quality control, the dataset comprised 32,163 genes and 32,842 spots across all samples, with comparable numbers of detected genes and sequencing counts across samples (Supplementary Figure 1a-b).

**Figure 1.**
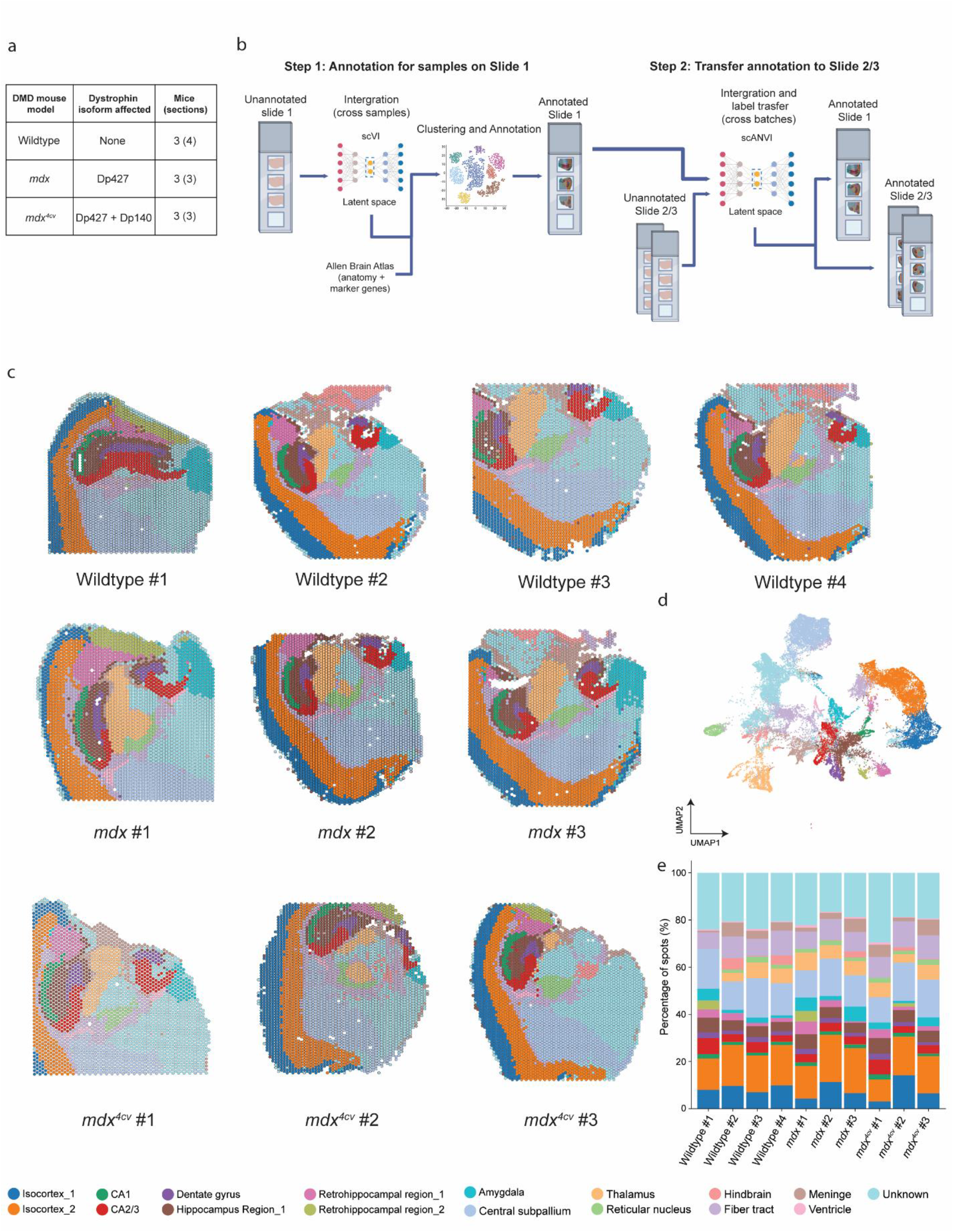
Annotation of the spatial transcriptomics dataset. (a) Overview of the DMD mouse models, dystrophin isoforms affected, numbers of mice, and Visium sections analyzed in this study. (b) Schematic overview of the annotation pipeline used in this study. Image by Biorender. (c) Spatial clustering results across all samples. (d) UMAP visualization of all spots in the integrated dataset, colored by brain region annotation. (e) Stacked bar plot showing the percentage of spots assigned to each cluster across samples.

In total, we identified 16 distinct brain regions, including multiple isocortical layers, the hippocampus, the amygdala and the hypothalamus (Figure 1c-d; Supplementary Figure 1c). However, because of anatomical complexity and ambiguity, we could not annotate all clusters, leading to one cluster being annotated as “unknown”. Visual comparison of brain-region composition across samples revealed that the distribution of brain regions was consistent across the dataset (Figure 1c,e).

### Broader transcriptomic alterations across brain regions following Dp140 loss

We sought identify genes associated with the loss of dystrophin isoforms across brain regions and mouse models. To distinguish transcriptional changes associated with the loss of different dystrophin isoforms, we used three genotype comparisons: WT and *mdx* brains were compared to identify transcriptional changes associated with the loss of the full-length Dp427 isoforms. WT and *mdx^4cv^* samples were compared to identify transcriptional changes associated with the loss of Dp427 and Dp140. Finally, *mdx* and *mdx^4cv^* brains were compared to identify changes associated with the loss of Dp140 on a Dp427 deficient background.

The additional loss of Dp140 was associated with broader transcriptional changes than the loss of Dp427 alone, as reflected by the higher number of DEGs identified in the WT versus *mdx^4cv^* comparison than in the WT versus *mdx* comparison. These broader changes were observed across several brain regions, including the thalamus, amygdala, isocortex, and hippocampus (Figure 2a, Supplementary Table 1-3).

**Figure 2.**
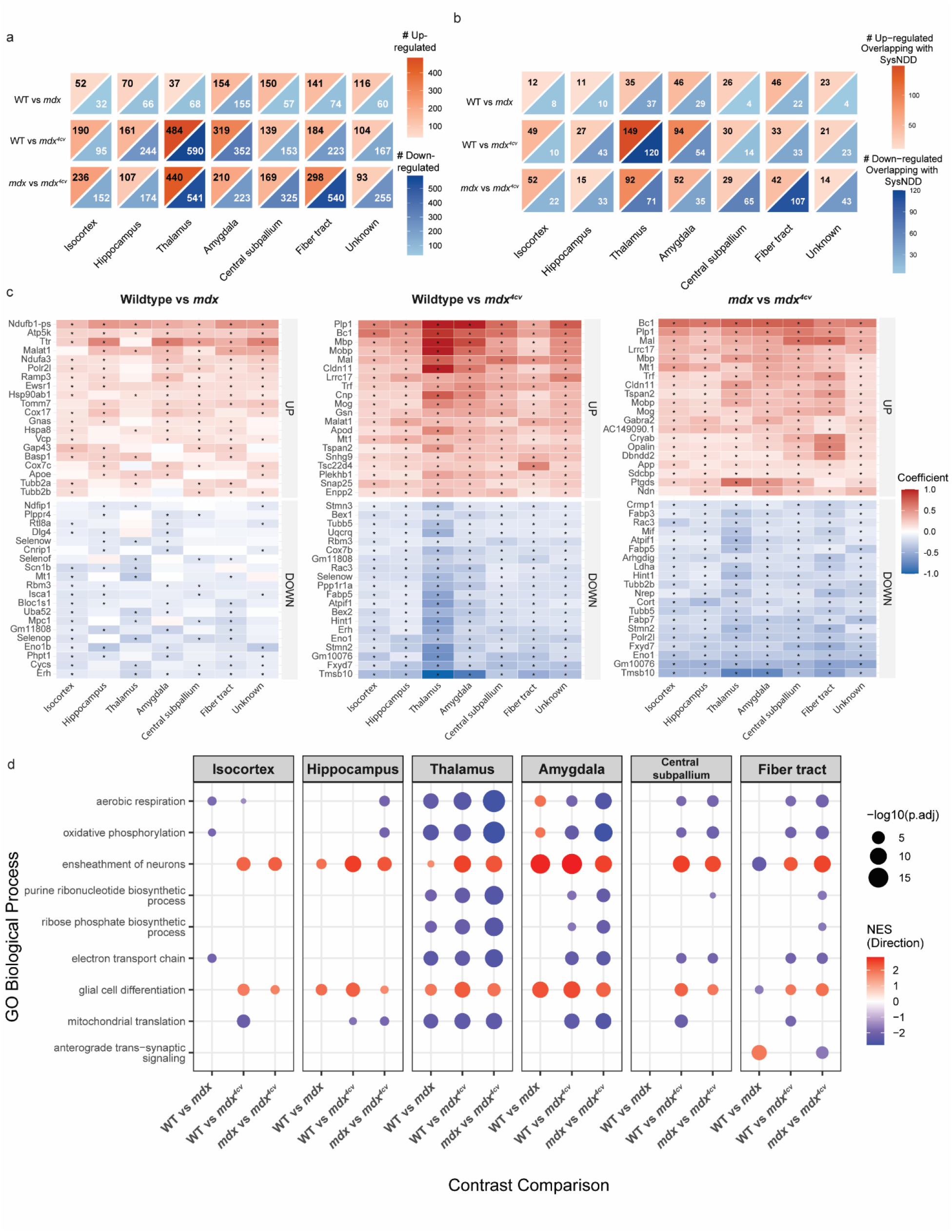
Differential gene expression associated with Dp427 and Dp140 deficiency across brain regions. (a) Triangular heatmap showing the number of upregulated and downregulated DEGs across high-level annotated brain regions. Heatmap colors indicate the direction of differential expression, with red representing upregulated genes and blue representing downregulated genes. (b) Triangular heatmap showing overlap between genes in the SysNDD database and region-specific upregulated and downregulated DEGs. Heatmap colors indicate the direction of differential expression, with red representing upregulated genes and blue representing downregulated genes. (c) Heatmap of top differentially expressed genes across ≥3 brain regions. Genes were ranked by the number of regions in which they were significantly differentially expressed, followed by their mean absolute MAST hurdle model coefficient. Asterisks denote significant differential expression within individual brain regions (|coefficient| > 0.1, Benjamini–Hochberg-adjusted P < 0.05). Colors represent the MAST hurdle model coefficient, with red indicating higher expression and blue indicating lower expression. (d) Top 5 enriched GO terms per genotype comparison from each brain region were combined across genotype comparisons. Redundant terms were consolidated using a Wang semantic similarity threshold of <0.6. Dot color represents the normalized enrichment score (NES), with red indicating positive enrichment and blue indicating negative enrichment, while dot size represents the adjusted P value. Only significantly enriched terms (FDR < 0.05) are shown; blank entries indicate non-significant enrichment.

Based on the symptoms observed in patients with DMD, next, we investigated whether the identified DEGs were enriched for genes associated with neurodevelopmental disorders (NDDs). To assess the relevance of our findings to NDDs, we examined the overlap between our DEG sets and NDD-associated genes curated in the SysNDD database (27) (Figure 2b). In *mdx* mice, we observed limited overlap between the DEGs and SysNDD genes, with the overlapping genes distributed relatively evenly across brain regions. In *mdx^4cv^*mice, the majority of overlapping genes were found in the thalamus, isocortex and hippocampus region, suggesting that the additional loss of Dp140 may particularly affect NDD-related genes in these brain regions.

We further ranked the top DEGs according to the breadth of their effects across brain regions and the magnitude of their strongest up or downregulation. This allowed us to identify transcriptional changes with broad effects across brain regions in different genotypes. The top-tier DEGs identified in *mdx^4cv^* mice showed greater magnitudes of upregulation and downregulation than those found in mdx mice, indicating a stronger and more widespread transcriptomic perturbation associated with the additional loss of Dp140 in *mdx^4cv^* mice (Figure 2c–d). Consistent with this finding, the direct comparison between *mdx* and *mdx^4cv^* mice revealed a set of DEGs that overlapped with those identified in the WT versus *mdx^4cv^* comparison, highlighting the additional impact of Dp140 loss on brain gene expression (Supplementary Figure 2a-b).

Specifically, we identified several mitochondrial respiration-related genes among the top DEGs, including *Atp5k*, *Ndufa3*, and *Cox7c*, which were consistently upregulated across multiple brain regions in *mdx* mice (Figure 2c). Gene set enrichment analysis (GSEA) identified mitochondrial respiration-related processes, including “aerobic respiration” and “oxidative phosphorylation”, among the most significantly enriched pathways (Figure 2d, Supplementary Table 4-6).

Mitochondrial pathway enrichment exhibited regional heterogeneity in *mdx* mice (Figure 2c). Although several mitochondrial genes were consistently upregulated across multiple brain regions in *mdx* mice, mitochondrial respiration gene sets were specifically downregulated in the isocortex and thalamus. as indicated by negative normalized enrichment scores (NESs). Fewer mitochondrial respiration-related genes were found among the top DEGs in *mdx^4cv^*mice when compared with either WT or *mdx* mice (Figure 2c). Nevertheless, in the *mdx* versus *mdx^4cv^*comparison, GSEA revealed negative NESs for mitochondrial respiration-related terms across all examined regions except the isocortex. This suggests that the loss of Dp140 induces subtle but coordinated changes across a broad set of mitochondrial genes, which are more readily captured by pathway-level analysis than by individual genes.

We also observed genes associated with myelination in the top DEG list, including *Mbp*, *Mobp*, *Mal*, and *Mog.* These genes were consistently upregulated across brain regions in *mdx^4cv^* mice compared to both WT and *mdx* mice. This indicates that the upregulation is specific to Dp140 loss, as it does not occur in *mdx* mice lacking only Dp427 (Figure 2c). Consistent with these gene expression patterns, GSEA analysis identified “ensheathment of neurons”, and “glial cell differentiation” as the predominant pathways especially in WT versus *mdx^4cv^* (Figure 2d). We also observed greater regional heterogeneity in *mdx* mice than in WT mice. Myelination associated gene sets were positively enriched in most of the brain regions but negatively enriched in the fiber tract, indicating region-specific regulation of myelination pathways. Overall, the WT versus *mdx^4cv^*comparison revealed widespread, significant positive enrichment of oligodendrocyte and myelin-associated pathways, indicating that the additional loss of Dp140 in *mdx^4cv^* mice leads to pronounced upregulation of these transcriptional programs.

### Enhanced myelination related gene expression in the thalamus following Dp140 loss

Building on the GSEA results highlighting consistent alterations in myelination and neuronal ensheathment across most regions, except for the fiber tracts, we sought to identify how dystrophin isoform loss affects the myelination-related gene programs across brain regions by focusing on the expression of genes mapped to "axon ensheathment". In *mdx* mice, this gene set was positively but relatively modestly enriched in most regions, with the notable exception of the fiber tracts, whereas in *mdx^4cv^*mice it was positively enriched in all regions examined (Figure 3a). This indicates that loss of Dp427 may be sufficient to modestly affect these pathways in multiple brain regions, but with a distinct direction of regulation in the fiber tracts. In contrast, the additional loss of Dp140 in *mdx^4cv^* mice may lead to broader and stronger pathway-level alterations across all brain regions.

**Figure 3.**
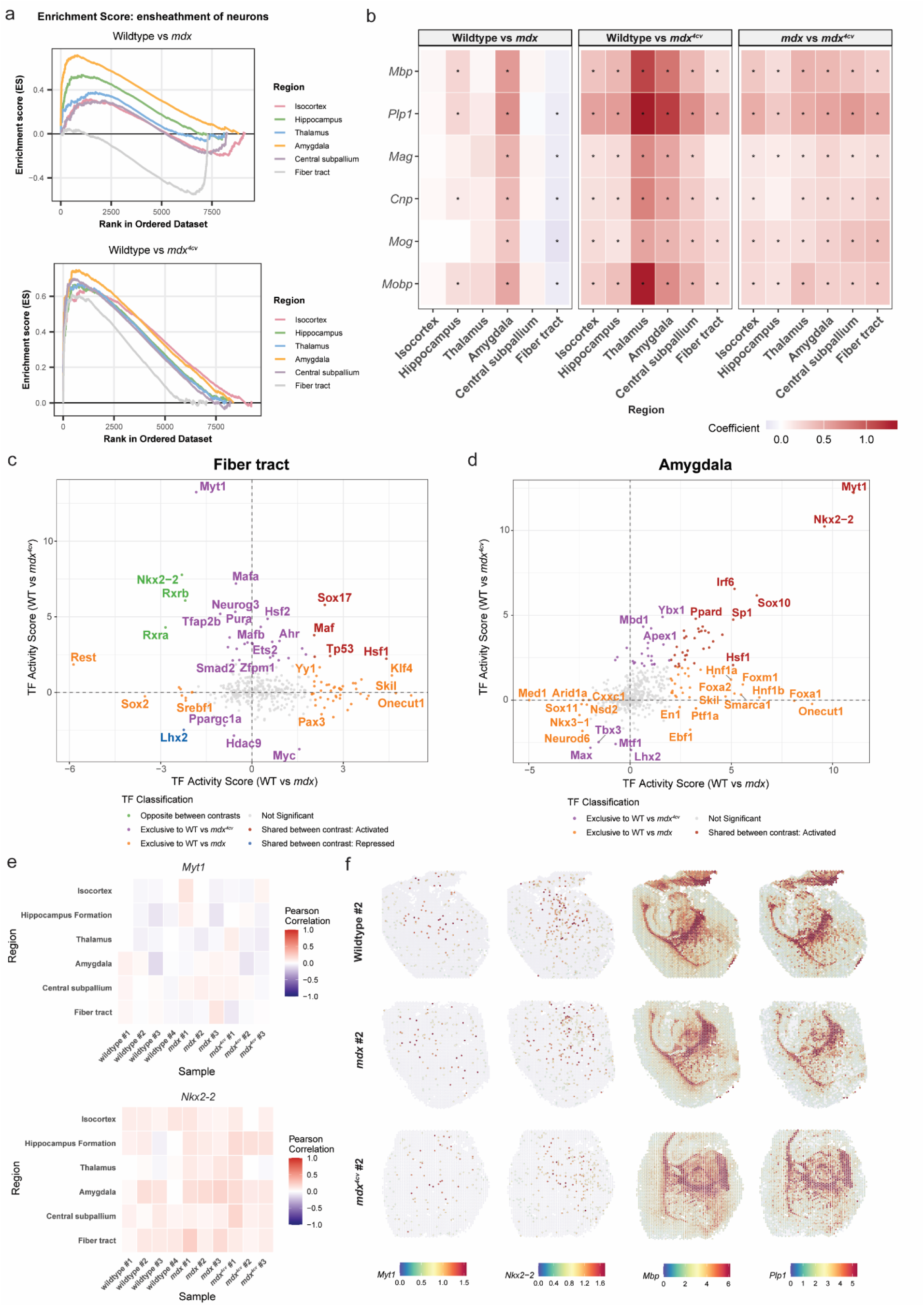
Dystrophin isoform deficiency disrupts myelination related transcriptional programs across brain regions. (a) Gene set enrichment analysis for neuronal ensheathment pathway across brain regions in (top) WT versus *mdx* and (bottom) WT versus *mdx^4cv^* comparisons. (b) Regional expression patterns of myelination markers across genotypes. Red and blue indicate positive and negative MAST hurdle model coefficients (representing the estimated expression effect size), respectively. Asterisks represent significant differential expression within individual regions (|coefficient| > 0.1 and Benjamini–Hochberg-adjusted *P* < 0.05). (c-d) Comparison of transcription factor activity scores between WT versus *mdx* and WT versus *mdx^4cv^* in the (c) fiber tract and (d) amygdala. Label colors classify TFs as showing opposite activity differences between contrasts (green), changes exclusive to the WT versus *mdx⁴ᶜᵛ* comparison (purple), changes exclusive to the WT versus *mdx* comparison (orange), or shared activity differences in the same direction in both comparisons (red: activated; blue: depressed). Activity scores estimate TF function based on the expression of known target genes. (e) Pearson correlations of regional coexpression patterns between log normalized expression of myelination genes and (top) *Myt1* and (bottom) *Nkx2-2* across biological samples. (f) Spatial expression of *Myt1*, *Nkx2-2*, *Mbp*, and *Plp1* in representative WT, *mdx*, and *mdx^4cv^* brain sections. Color intensity indicates the log normalized expression level of each gene.

**Figure 4.**
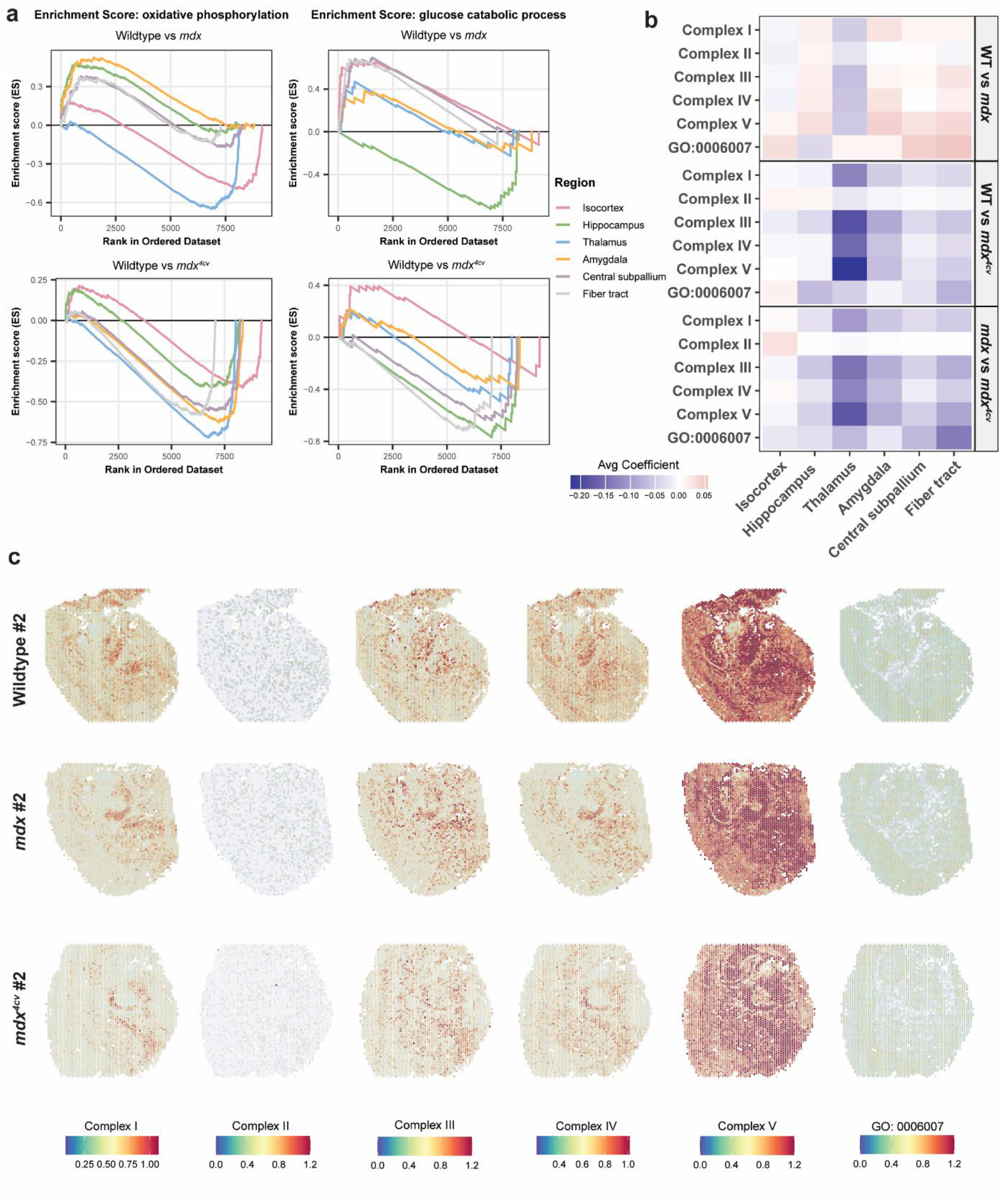
Brain region specific alterations in oxidative phosphorylation and glucose catabolism upon dystrophin isoform deficiency. (a) Gene set enrichment analysis of oxidative phosphorylation (left) and glucose catabolic process (right) across brain regions in WT versus *mdx* (top) and WT versus *mdx^4cv^* (bottom) comparisons. (b) Heatmap showing the region average differential expression coefficients of specific gene sets associated with mitochondrial respiratory chain complexes I–V and the glucose catabolic process (GO:0006007) in brain regions, separated by genotype comparison. Color gradients indicate pathway-level upregulation (red) or downregulation (blue). (c) Spatial distribution of module scores for gene sets associated with mitochondrial respiratory chain complexes I–V and the glucose catabolic process (GO:0006007) in representative WT, *mdx*, and *mdx^4cv^* brain sections.

Next, we selected six myelination associated markers from previous studies (*Mbp*, *Plp1*, *Mag*, *Cnp*, *Mog*, *Mobp*). We assessed their expression changes across brain regions and genotypes to determine whether the gene-level patterns were consistent with the myelination-related pathway changes identified by GSEA (31-33). We observed that these myelination associated genes were mainly expressed in thalamus, reticular nucleus, central subpallium and fiber tract (Supplementary Figure 3a-f). Expression changes in these myelination markers were significantly divergent across regions, mirroring the pathway-level trends. In the WT versus *mdx* comparison, they were downregulated in the fiber tracts but showed little or weak upregulation across other brain regions, with relatively stronger upregulation in the amygdala (Figure 3b). The six myelination markers were expressed at higher levels in *mdx^4cv^*, when compared to WT and *mdx* mice (Figure 2b), indicating that additional loss of Dp140 is associated with a stronger shift in myelination program than loss of Dp427 alone. This finding also suggests that the fiber tract are differentially sensitive to the loss of Dp427 and/or Dp140 in our study.

To identify the upstream transcriptional regulators driving the observed gene expression changes, we used decoupleR to infer transcription factor (TF) activity based on the differential expression of known target genes across brain regions (Figure 3c–d). In the fiber tracts, both *Myt1* and *Nkx2-2* were downregulated in *mdx* mice but upregulated in *mdx^4cv^* mice relative to WT mice, indicating that the additional loss of Dp140 not only overrides but also reverses the repression caused by Dp427 loss. In contrast, in the amygdala, these factors showed enhanced expression in both DMD models, indicating that the loss of Dp427 alone drives the response and that the additional loss of Dp140 contributes to a lesser extent. Together, these results indicate that the enhancement or suppression of myelination-related TF activity caused by dystrophin deficiency depends on the brain region and the pattern of isoform loss.

Correlation analysis between the expression of *Myt1* and *Nkx2-2* and the myelination associated genes yielded weak associations (Figure 3e–f), consistent with their weak and sparse expression. Together, these results reveal widespread but regionally heterogeneous changes in myelination associated transcriptional and regulatory programs in the DMD murine brain, with the most pronounced changes in the *mdx^4cv^* model.

### Dystrophin deficiency is associated with brain region specific alterations in energy metabolism

Next, we focused on two representative energy metabolism terms drawn from the pathways associated with the top DEGs: "oxidative phosphorylation" and "glucose catabolic process"(GO:0006007) (Supplementary Figure 4). The loss of full-length Dp427 in *mdx* mice was associated with bidirectional, region-specific expression shifts: GSEA of "oxidative phosphorylation" revealed positive enrichment in the amygdala and negative enrichment in the thalamus and isocortex, along with negative enrichment of “glucose metabolic process” in the hippocampus. On the other hand, we observed that the additional loss of Dp140 in the *mdx^4cv^* model corresponded to a broader suppression of energy metabolism pathways. In the WT versus *mdx^4cv^* comparison, "oxidative phosphorylation" was negatively enriched across thalamus, amygdala, central subpallium, and fiber tract. Additionally, "glucose catabolic process" was negatively enriched in the hippocampus in the WT versus *mdx^4cv^* comparison, and across the thalamus, amygdala, central subpallium, and fiber tracts in the *mdx* versus *mdx^4cv^* comparison. Together, these findings indicate a widespread dysregulation of energy metabolism of *mdx^4cv^* mice due to loss of Dp140.

To resolve these pathway-level trends in greater detail, we examined gene signatures representing each mitochondrial respiratory chain complex, curated from the expert-annotated MitoCarta3.0 database, excluding mitochondrially encoded (*Mt-*) genes from the analysis (Supplementary Table 7) (29). Consistent with the pathway level findings, signatures for respiratory chain complexes I–V and glucose catabolism showed heterogeneous changes associated with the isolated loss of Dp427. In *mdx* mice, we observed a general upregulation of electron transport chain (ETC) gene expression across most brain regions, with the notable exception of the thalamus and isocortex. In contrast, the loss of Dp427 and Dp140 resulted in broad downregulation across these signatures. In *mdx^4cv^* mice, the thalamus showed a pronounced reduction in the expression of all five respiratory chain complexes. We also examined region specific genes under the “glucose metabolic process” term. Associated with Dp427 loss in *mdx*, we found glucose catabolism was negatively enriched in the hippocampus, whereas the additional loss of shorter isoforms in *mdx^4cv^* predominantly corresponded to a broader downregulation, except for the isocortex.

We next mapped the spatial expression pattern of oxidative phosphorylation complex and glucose metabolic process pathway across WT, *mdx*, and *mdx^4cv^* brain sections (Figure 3c, Supplementary Figure 5a-f). Complexes I, III, IV, and V showed comparatively higher expression, whereas Complex II appeared sparse, likely reflecting the smaller number of genes represented in this signature. Across all electron transport chain complexes, expression was consistently high in the reticular nucleus of the thalamus and in the central subpallium, and low in the outer cortical layers and dentate gyrus. The glucose metabolic process pathway, by contrast, was most highly expressed in the inner cortical layers, CA2/3, amygdala, and amygdala but low expressed in fiber tract.

## Discussion

We applied spatial transcriptomics to systematically profile the transcriptional alterations resulting from the loss of full-length Dp427 isoforms and the additional shorter isoform Dp140 across multiple brain regions in *mdx* and *mdx^4cv^* mice in the early postnatal stage. This spatially-resolved approach allowed us to characterize global transcriptional alterations, while also identifying region-specific molecular signatures associated with dystrophin isoform deficiency. Moreover, by incorporating the *mdx^4cv^*model, our work addresses an important gap in knowledge by providing whole transcriptome insight into the impact of Dp140 deficiency across brain regions, as the function of Dp140 remains underexplored relative to the more widely studied full-length Dp427 isoforms.

We found that the severity and pattern of transcriptional dysregulation depends on which specific dystrophin isoforms were deficient, as well as the brain region examined. The *mdx^4cv^* model showed more differentially expressed genes than the *mdx* model, suggesting that the additional loss of Dp140 expands the transcriptional consequences of dystrophin deficiency. This is in line with clinical findings showing that mutations affecting distal dystrophin isoforms are often linked to more pronounced cognitive impairment (34, 35). Notably, these changes were not evenly distributed across the brain: DEGs between WT and *mdx^4cv^* mice were enriched in the thalamus, amygdala, isocortex, and hippocampus, pointing to these regions as particularly vulnerable to Dp140 isoform loss. Furthermore, particularly in the thalamus, amygdala, and isocortex, there was substantial overlap with genes associated with neurodevelopment disorders, suggesting that these alterations are linked to cognitive disorders caused by Dp140 deficiency.

Expression of Dp140 is an isoform whose brain expression is developmentally regulated. Dp140 is predominantly expressed in the developing human brain, including the telencephalic cortical plate, dorsal hippocampus and thalamic neuroepithelium, before its expression is markedly reduced in the adult brain (15). Similar patterns have been observed in the mouse cortex and cerebellum where Dp140 protein levels are more abundant at P14 than at later developmental stages. This suggests that Dp140 might be important for synaptogenesis and glutamatergic neurotransmission during this developmental window (36). The functional relevance of Dp140 loss is further supported by studies in *mdx52* mice which, similar to the *mdx^4cv^* model used in our study, lack both Dp427 and Dp140. *Mdx52* mice show reduced glutamatergic transmission in the basolateral amygdala alongside ASD-like behavior evident already in pups, and both electrophysiological and behavioral phenotypes are partially ameliorated by Dp140 restoration (37).

By examining DEGs across different regions and genotypes, we found transcriptional changes related to myelin/oligodendrocytes related pathways in both *mdx* and *mdx^4cv^*brain samples, though the two models showed different patterns. In *mdx* mice, these markers were upregulated in the amygdala and hippocampus, but downregulated in fiber tract regions. Aranmolate *et al.* established that oligodendrocytes express three dystrophin isoforms (Dp427, Dp140 and Dp71), with expression levels increasing as OPCs differentiate into mature oligodendrocytes (38). They also reported that *mdx* mice have reduced myelin basic protein (MBP) in cortical lysates during early postnatal myelination (P14–P42), pointing to developmentally delayed cortical myelination (38). Another study further showed age-dependent disturbances of postnatal oligodendrogenesis in the *mdx* mouse, with fewer mature oligodendrocytes in the corpus callosum at postnatal day 21, and transmission electron microscopy revealing thinner myelin, delayed myelin compaction and abnormal myelin structures that persisted into adulthood (39). The reduction in corpus callosum MBP at P14 in their data aligns with the reduced myelin and oligodendrocyte transcript signals we observed in *mdx* fiber tract regions at P14. However, CNP and MOG protein levels were largely unchanged at P14 in that study. This suggests that the transcript level perturbations we detected are unlikely to be mirrored consistently across all myelin and oligodendrocyte protein markers, and that the timing of transcriptional and protein changes may not be fully aligned.

Neuroimaging studies also provide evidence of impaired myelination in DMD patients. Diffusion tensor imaging in DMD patients revealed white matter microstructural abnormalities. Doorenweerd *et al.* reported reduced fractional anisotropy (FA) in the occipital lobe, higher mean diffusivity in the corpus callosum, and higher radial diffusivity throughout the white matter in boys with DMD (40). Fu *et al.* found reduced splenium FA correlating with lower verbal IQ (41). Since elevated radial diffusivity is a recognized correlate of demyelination, though reduced FA may also reflect axonal density or membrane changes, these signatures echo with the delayed, abnormal myelination seen in the *mdx* corpus callosum, bridging our transcriptional changes and the DMD cognitive phenotype.

We also observed transcriptional changes related to oxidative phosphorylation in both *mdx* and *mdx^4cv^*brain samples but with different patterns. The isolated loss of Dp427 resulted in an upregulation of ETC genes across most detected brain regions, accompanied by regional downregulation in the thalamus and isocortex. In contrast, the loss of Dp427 and Dp140 drove a widespread downregulation of ETC genes across nearly all brain regions, with the most pronounced reduction localized to the thalamus. In skeletal muscle, dystrophin deficiency is known to cause oxidative stress, calcium overload, and early mitochondrial damage (42-44).

Measurements of brain mitochondrial function in *mdx* mice have shown different results across studies. Pomeroy *et al.* revealed upregulation of nuclear encoded ETC subunits in cerebra of 10 day old *mdx* mice, alongside a decrease in mitochondrially encoded transcripts (16). In our data, nuclear-encoded ETC genes were upregulated in most of the detected regions except for thalamus and isocortex, indicating that whole cerebrum measurements may average over opposing regional trends. Bellissimo *et al.* found that 4-week-old D2-*mdx* mice (lacking Dp427) had reduced hippocampal Complex I-supported respiration relative to wild type mice, which was restored by treatment with the adiponectin receptor agonist ALY688 (45). In addition, Tuon *et al.* measured ETC enzyme activities in 3-month-old *mdx* brains and reported reduced Complex I activity across multiple regions and reduced Complex IV activity in prefrontal cortex and striatum (46). The differences likely reflect variation in mouse models, developmental stage, brain regions included, and whether abundance was measured at the protein or transcript level. Nonetheless, they collectively indicate that brain mitochondrial dysfunction is a recurring feature from early postnatal life through adulthood in *mdx* mice.

We also noticed a broad downregulation of ETC genes across brain regions due to additional loss of Dp140 in *mdx^4cv^* mice, which has not been previously reported to our knowledge. Murphy *et al.* performed label-free proteomics in 12-month-old *mdx^4cv^* brains and observed altered abundance of proteins involved in energy metabolism, ion handling, and glial structure indicating metabolic adaptation and disturbed ion homeostasis (47). Direct comparison with our data is limited by different developmental stages and modalities. Reduced mitochondrial gene expression is also being observed in autism post-mortem brain samples (48-50), and schizophrenia post-mortem prefrontal cortex (51, 52). The broader reduction of mitochondrial genes in *mdx^4cv^* mice is consistent with the clinical picture in which loss of Dp140 alongside Dp427 in DMD patients is associated with a more severe cognitive and neurodevelopmental phenotype, including lower IQ, a higher prevalence of intellectual disability, and clinically reported autism spectrum features.

Several limitations are warranted when interpreting our findings. Firstly, our spatial data measures gene expression levels, and changes in mRNA do not always translate into corresponding changes in protein or function. Several transcriptomic changes identified in the myelin and oxidative phosphorylation pathways were not consistent with previously reported protein and enzyme activity data. This likely reflects post-transcriptional or compensatory regulatory mechanisms, highlighting that mRNA changes do not necessarily translate into corresponding protein-level effects. Establishing the functional relevance of these transcriptomic alterations will require region-specific protein and functional analyses in the same models as used in this study. Secondly, our analyses capture a single developmental stage, which limits our ability to track the timing of these changes. Although cortical Dp140 is largely downregulated after birth in humans, it remains present in the postnatal mouse brain (14, 53). Thus, while our timepoint successfully captures ongoing Dp140 expression, a full time course spanning fetal, early and late postnatal stages is still required to characterize the temporal dynamics of transcriptional changes following Dp140 loss. Thirdly, although Visium preserves regional context, each spot contains several cells, leaving our region level signals unassigned to specific cell populations. Finally, our findings are based on mouse models and a limited sample size, so further exploration to the human DMD brain remains to be established and the neuroimaging parallels we draw point in a consistent direction but without being conclusive.

In summary, the loss of dystrophin produces distinct transcriptional changes in the brain that vary by region rather than affecting all areas equally, converging on myelin and oligodendrocyte biology and oxidative phosphorylation. These changes depend on which dystrophin isoforms are absent and differ between the *mdx* and *mdx^4cv^* models. Their pattern corresponds with the greater severity of cognitive impairment reported in patients with distal mutations. The molecular basis this work provides could facilitate future work to define these mechanisms and explore therapeutic targets.

## CRediT authorship contribution statement

**Qirong Mao**: Conceptualization, Data curation, Formal analysis, Methodology, Software, Visualization, Writing – original draft, Writing – review and editing. **Laura G.M. Heezen**: Resources, Investigation, Writing – review and editing. **Sharon de Vries**: Resources, Investigation, Writing – review and editing. **Nynke Verwoerd**: Investigation. **Annemieke Aartsma-Rus**: Supervision, Writing – review and editing. **Maaike van Putten**: Conceptualization, Resources, Supervision, Writing – review and editing. **Pietro Spitali**: Conceptualization, Methodology, Funding acquisition, Project administration, Resources, Supervision, Writing – review and editing. **Ahmed Mahfouz**: Conceptualization, Methodology, Project administration, Supervision, Writing – review and editing.

## Acknowledgement

We acknowledge funding by the Medical Genomics Theme and the Human Genetics Department of the LUMC. We would like to thank Tiberiu Stan, Christa Tanganyika-de Winter (Leiden University Medical Center; Leiden, The Netherlands) and Lizette van der Pijl for helping sample preprocessing. We also would like to thank Lola van Doorne for helping perform parts of the Visium experiments.

## Data availability

The spatial transcriptomics datasets generated in the current study have been deposited in the Gene Expression Omnibus (GEO). The dataset is currently private and will be made publicly available upon official publication of the associated manuscript.

## Code availability

Scripts used for the data analysis presented in this study are available at: https://github.com/Qirongmao97/Visium_brain.

## Reference

1. Mendell JR, Shilling C, Leslie ND, Flanigan KM, al-Dahhak R, Gastier-Foster J, et al. Evidence-Based Path to Newborn Screening for Duchenne Muscular Dystrophy. Ann Neurol. 2012;71(3):304–13.

2. Zhang XF, Luo YY, Jiang L, Hong SQ. Clinical study on cognitive impairment in Duchenne muscular dystrophy. Neuromuscul Disord. 2023;33(7):596–604.

3. Banihani R, Smile S, Yoon G, Dupuis A, Mosleh M, Snider A, et al. Cognitive and Neurobehavioral Profile in Boys With Duchenne Muscular Dystrophy. J Child Neurol. 2015;30(11):1472–82.

4. Cotton S, Voudouris NJ, Greenwood KM. Intelligence and Duchenne muscular dystrophy: full-scale, verbal, and performance intelligence quotients. Dev Med Child Neurol. 2001;43(7):497–501.

5. Darmahkasih AJ, Rybalsky I, Tian CX, Shellenbarger KC, Horn PS, Lambert JT, et al. Neurodevelopmental, behavioral, and emotional symptoms common in Duchenne muscular dystrophy. Muscle Nerve. 2020;61(4):466–74.

6. Pane M, Lombardo ME, Alfieri P, D’Amico A, Bianco F, Vasco G, et al. Attention Deficit Hyperactivity Disorder and Cognitive Function in Duchenne Muscular Dystrophy: Phenotype-Genotype Correlation. J Pediatr-Us. 2012;161(4):705-+.

7. Panda PK, Shikha D, Sharawat IK. Association of Autism Spectrum Disorder with Pre-symptomatic Duchenne Muscular Dystrophy: Isolated Elevation of Transaminases as a Diagnostic Clue. Ann Indian Acad Neur. 2021;24(4):603–4.

8. Lee AJ, Buckingham ET, Kauer AJ, Mathews KD. Descriptive Phenotype of Obsessive Compulsive Symptoms in Males With Duchenne Muscular Dystrophy. J Child Neurol. 2018;33(9):572–9.

9. Hinton VJ, De Vivo DC, Fee R, Goldstein E, Stern Y. Investigation of Poor Academic Achievement in Children with Duchenne Muscular Dystrophy. Learn Disabil Res Pract. 2004;19(3):146–54.

10. Miranda R, Weerkamp PMM, Kolesnik A, Geagan C, Chieffo D, Suarez-Bagnasco M, et al. Screening for brain-related comorbidities in Duchenne muscular dystrophy: Construction, reliability, and validity of the BIND screener. Dev Med Child Neurol. 2026;68(8):1080–96.

11. Ueda Y, Suwazono S, Maedo S, Higuchi I. Profile of cognitive function in adults with duchenne muscular dystrophy. Brain Dev-Jpn. 2017;39(3):225–30.

12. Tizzoni F, Canella G, Delle Fave A, Di Lernia D, Lorusso ML, Nobile M, et al. Living with Duchenne Muscular Dystrophy Beyond the Physical Implications: Cognitive Features, Psychopathology Aspects, and Psychosocial Resources-A Narrative Review. Brain Sciences. 2025;15(7).

13. Muntoni F, De Waele L, Munell F, Schara-Schmidt U, Mercuri E, Hendriksen J, et al. The unmet need of psychopharmacological intervention for neuropsychiatric and neurodevelopmental comorbidities in Duchenne muscular dystrophy: report of 5 workshops. Eur J Paediatr Neurol. 2026;61:84–9.

14. Doorenweerd N, Mahfouz A, van Putten M, Kaliyaperumal R, t’ Hoen PAC, Hendriksen JGM, et al. Timing and localization of human dystrophin isoform expression provide insights into the cognitive phenotype of Duchenne muscular dystrophy. Sci Rep-Uk. 2017;7.

15. Catapano F, Alkharji R, Chambers D, Singh S, Aghaeipour A, Malhotra J, et al. A comprehensive spatiotemporal map of dystrophin isoform expression in the developing and adult human brain. Acta Neuropathol Com. 2025;13(1).

16. Pomeroy J, Borczyk M, Kawalec M, Hajto J, Carlson E, Svärd S, et al. Spatiotemporal diversity in molecular and functional abnormalities in the mdx dystrophic brain. Molecular Medicine. 2025;31(1).

17. Saoudi A, Mitsogiannis MD, Zarrouki F, Fergus C, Stojek E, Talavera S, et al. Impact of distinct dystrophin gene mutations on behavioral phenotypes of Duchenne muscular dystrophy. Disease Models & Mechanisms. 2024;17(12).

18. Bulfield G, Siller WG, Wight PA, Moore KJ. X chromosome-linked muscular dystrophy (mdx) in the mouse. Proc Natl Acad Sci U S A. 1984;81(4):1189–92.

19. Im WB, Phelps SF, Copen EH, Adams EG, Slightom JL, Chamberlain JS. Differential expression of dystrophin isoforms in strains of mdx mice with different mutations. Hum Mol Genet. 1996;5(8):1149–53.

20. Ratz M, von Berlin L, Larsson L, Martin M, Westholm JO, La Manno G, et al. Clonal relations in the mouse brain revealed by single-cell and spatial transcriptomics. Nat Neurosci. 2022;25(3):285–94.

21. Wolf FA, Angerer P, Theis FJ. SCANPY: large-scale single-cell gene expression data analysis. Genome Biol. 2018;19(1):15.

22. Lopez R, Regier J, Cole MB, Jordan MI, Yosef N. Deep generative modeling for single-cell transcriptomics. Nat Methods. 2018;15(12):1053–8.

23. Hao Y, Stuart T, Kowalski MH, Choudhary S, Hoffman P, Hartman A, et al. Dictionary learning for integrative, multimodal and scalable single-cell analysis. Nat Biotechnol. 2024;42(2):293–304.

24. R Core Team. R: A Language and Environment for Statistical Computing. R Foundation for Statistical Computing; 2023.

25. Finak G, McDavid A, Yajima M, Deng J, Gersuk V, Shalek AK, et al. MAST: a flexible statistical framework for assessing transcriptional changes and characterizing heterogeneity in single-cell RNA sequencing data. Genome Biol. 2015;16:278.

26. Wu T, Hu E, Xu S, Chen M, Guo P, Dai Z, et al. clusterProfiler 4.0: A universal enrichment tool for interpreting omics data. Innovation (Camb). 2021;2(3):100141.

27. Kochinke K, Zweier C, Nijhof B, Fenckova M, Cizek P, Honti F, et al. Systematic Phenomics Analysis Deconvolutes Genes Mutated in Intellectual Disability into Biologically Coherent Modules. Am J Hum Genet. 2016;98(1):149–64.

28. Badia IMP, Velez Santiago J, Braunger J, Geiss C, Dimitrov D, Muller-Dott S, et al. decoupleR: ensemble of computational methods to infer biological activities from omics data. Bioinform Adv. 2022;2(1):vbac016.

29. Rath S, Sharma R, Gupta R, Ast T, Chan C, Durham TJ, et al. MitoCarta3.0: an updated mitochondrial proteome now with sub-organelle localization and pathway annotations. Nucleic Acids Research. 2021;49(D1):D1541–D7.

30. Sunkin SM, Ng L, Lau C, Dolbeare T, Gilbert TL, Thompson CL, et al. Allen Brain Atlas: an integrated spatio-temporal portal for exploring the central nervous system. Nucleic Acids Research. 2013;41(D1):D996–D1008.

31. Fulton D, Paez PM, Campagnoni AT. The multiple roles of myelin protein genes during the development of the oligodendrocyte. ASN Neuro. 2010;2(1):e00027.

32. Kaushansky N, Eisenstein M, Zilkha-Falb R, Ben-Nun A. The myelin-associated oligodendrocytic basic protein (MOBP) as a relevant primary target autoantigen in multiple sclerosis. Autoimmun Rev. 2010;9(4):233–6.

33. Johns TG, Bernard CCA. The structure and function of myelin oligodendrocyte glycoprotein. J Neurochem. 1999;72(1):1–9.

34. Felisari G, Boneschi FM, Bardoni A, Sironi M, Comi GP, Robotti M, et al. Loss of Dp140 dystrophin isoform and intellectual impairment in Duchenne dystrophy. Neurology. 2000;55(4):559–64.

35. Chamova T, Guergueltcheva V, Raycheva M, Todorov T, Genova J, Bichev S, et al. Association Between Loss of Dp140 and Cognitive Impairment in Duchenne and Becker Dystrophies. Balk J Med Genet. 2013;16(1):21–9.

36. Tetorou K, Aghaeipour A, Ma SY, Gileadi T, Saoudi A, Quinteiro PP, et al. Regional Expression of Dystrophin Gene Transcripts and Proteins in the Mouse Brain. Cells. 2025;14(18).

37. Hashimoto Y, Kuniishi H, Sakai K, Fukushima Y, Du X, Yamashiro K, et al. Brain Dp140 alters glutamatergic transmission and social behaviour in the mouse model of Duchenne muscular dystrophy. Progress in Neurobiology. 2022;216.

38. Aranmolate A, Tse N, Colognato H. Myelination is delayed during postnatal brain development in the mouse model of Duchenne muscular dystrophy. Bmc Neurosci. 2017;18.

39. Arreguin AJ, Shao Z, Colognato H. Dmd mdx mice have defective oligodendrogenesis, delayed myelin compaction and persistent hypomyelination. Dis Model Mech. 2024;17(4).

40. Doorenweerd N, Straathof CS, Dumas EM, Spitali P, Ginjaar IB, Wokke BH, et al. Reduced Cerebral Gray Matter and Altered White Matter in Boys with Duchenne Muscular Dystrophy. Ann Neurol. 2014;76(3):403–11.

41. Fu Y, Dong YR, Zhang C, Sun Y, Zhang S, Mu XT, et al. Diffusion tensor imaging study in Duchenne muscular dystrophy. Ann Transl Med. 2016;4(6).

42. Moore TM, Lin AJ, Strumwasser AR, Cory K, Whitney K, Ho T, et al. Mitochondrial Dysfunction Is an Early Consequence of Partial or Complete Dystrophin Loss in Mice. Frontiers in Physiology. 2020;11.

43. Mareedu S, Million ED, Duan D, Babu GJ. Abnormal Calcium Handling in Duchenne Muscular Dystrophy: Mechanisms and Potential Therapies. Front Physiol. 2021;12:647010.

44. Bellissimo CA, Garibotti MC, Perry CGR. Mitochondrial stress responses in Duchenne muscular dystrophy: metabolic dysfunction or adaptive reprogramming? Am J Physiol Cell Physiol. 2022;323(3):C718–C30.

45. Bellissimo CA, Castellani LN, Finch MS, Murugathasan M, Gandhi S, Sweeney G, et al. Memory impairment in the D2.*mdx* mouse model of Duchenne muscular dystrophy is prevented by the adiponectin receptor agonist ALY688. Exp Physiol. 2023;108(9):1108–17.

46. Tuon L, Comim CM, Fraga DB, Scaini G, Rezin GT, Baptista BR, et al. Mitochondrial Respiratory Chain and Creatine Kinase Activities in Mouse Brain. Muscle Nerve. 2010;41(2):257–60.

47. Murphy S, Zweyert M, Henry M, Meleady P, Mundegar RR, Swandulla D, et al. Label-free mass spectrometric analysis reveals complex changes in the brain proteome from the *mdx-4cv* mouse model of Duchenne muscular dystrophy. Clin Proteom. 2015;12.

48. Anitha A, Nakamura K, Thanseem I, Matsuzaki H, Miyachi T, Tsujii M, et al. Downregulation of the Expression of Mitochondrial Electron Transport Complex Genes in Autism Brains. Brain Pathol. 2013;23(3):294–302.

49. Ginsberg MR, Rubin RA, Falcone T, Ting AH, Natowicz MR. Brain Transcriptional and Epigenetic Associations with Autism. Plos One. 2012;7(9).

50. Schwede M, Nagpal S, Gandal MJ, Parikshak NN, Mirnics K, Geschwind DH, et al. Strong correlation of downregulated genes related to synaptic transmission and mitochondria in post-mortem autism cerebral cortex. J Neurodev Disord. 2018;10.

51. Arion D, Corradi JP, Tang S, Datta D, Boothe F, He A, et al. Distinctive transcriptome alterations of prefrontal pyramidal neurons in schizophrenia and schizoaffective disorder. Mol Psychiatr. 2015;20(11):1397–405.

52. Prabakaran S, Swatton JE, Ryan MM, Huffaker SJ, Huang JTJ, Griffin JL, et al. Mitochondrial dysfunction in schizophrenia: evidence for compromised brain metabolism and oxidative stress. Mol Psychiatr. 2004;9(7):684–97.

53. Tetorou K, Aghaeipour A, Ma S, Gileadi T, Saoudi A, Perdomo Quinteiro P, et al. Regional Expression of Dystrophin Gene Transcripts and Proteins in the Mouse Brain. Cells. 2025;14(18).

