## Supplementary Figures for "Spatial transcriptomics maps region and isoform specific brain alterations in *mdx* and *mdx^4cv^* models of Duchenne muscular dystrophy"

**Supplementary Tables:**

**Supplementary Table 1.** Differentially expressed genes across brain regions in Wildtype versus *mdx* mice.

**Supplementary Table 2.** Differentially expressed genes across brain regions in Wildtype versus *mdx*^4cv^ mice.

**Supplementary Table 3.** Differentially expressed genes across brain regions in *mdx* versus *mdx*^4cv^ mice.

**Supplementary Table 4.** Region-specific GSEA of differential gene expression in Wildtype versus mdx mouse brains.

**Supplementary Table 5.** Region-specific GSEA of differential gene expression in Wildtype versus *mdx*^4cv^ mouse brains.

**Supplementary Table 6.** Region-specific GSEA of differential gene expression in *mdx* versus *mdx*^4cv^ mouse brains.

**Supplementary Table 7.** Region-specific coefficients of mitochondrial respiratory chain complexes I–V genes derived from MitoCarta 3.0 excluding mitochondrial genome-encoded (*MT-*) genes, and the glucose catabolic process (GO:0006007).

**Supplementary Figures:**

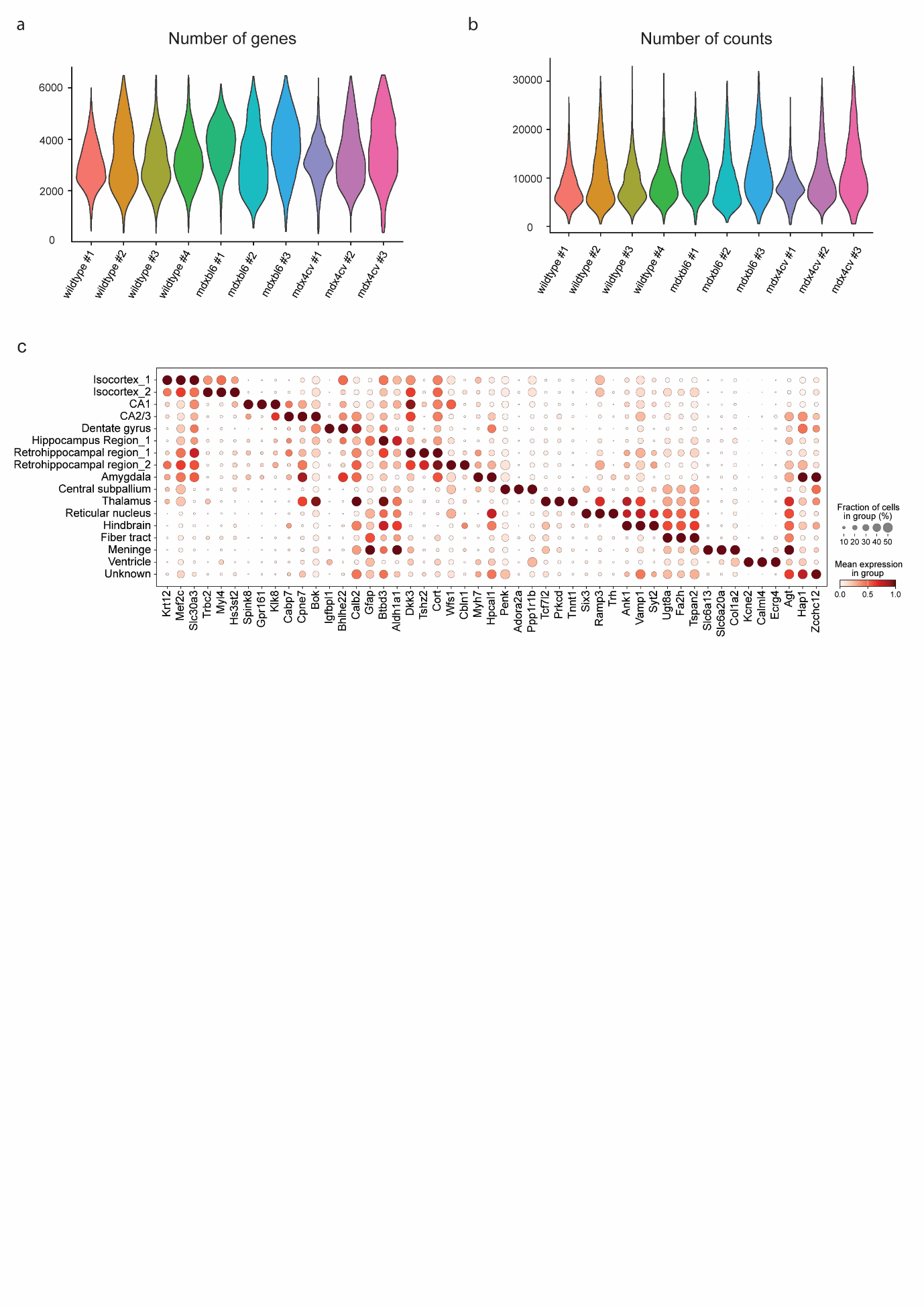

**Supplementary Figure 1. Quality control metrics and cell-type annotation of spatial transcriptomic datasets**

(a) Violin plot of numbers of genes per sample

(b) Violin plot of numbers of counts per sample

(c) Dot plot of marker gene expression across annotated brain regions

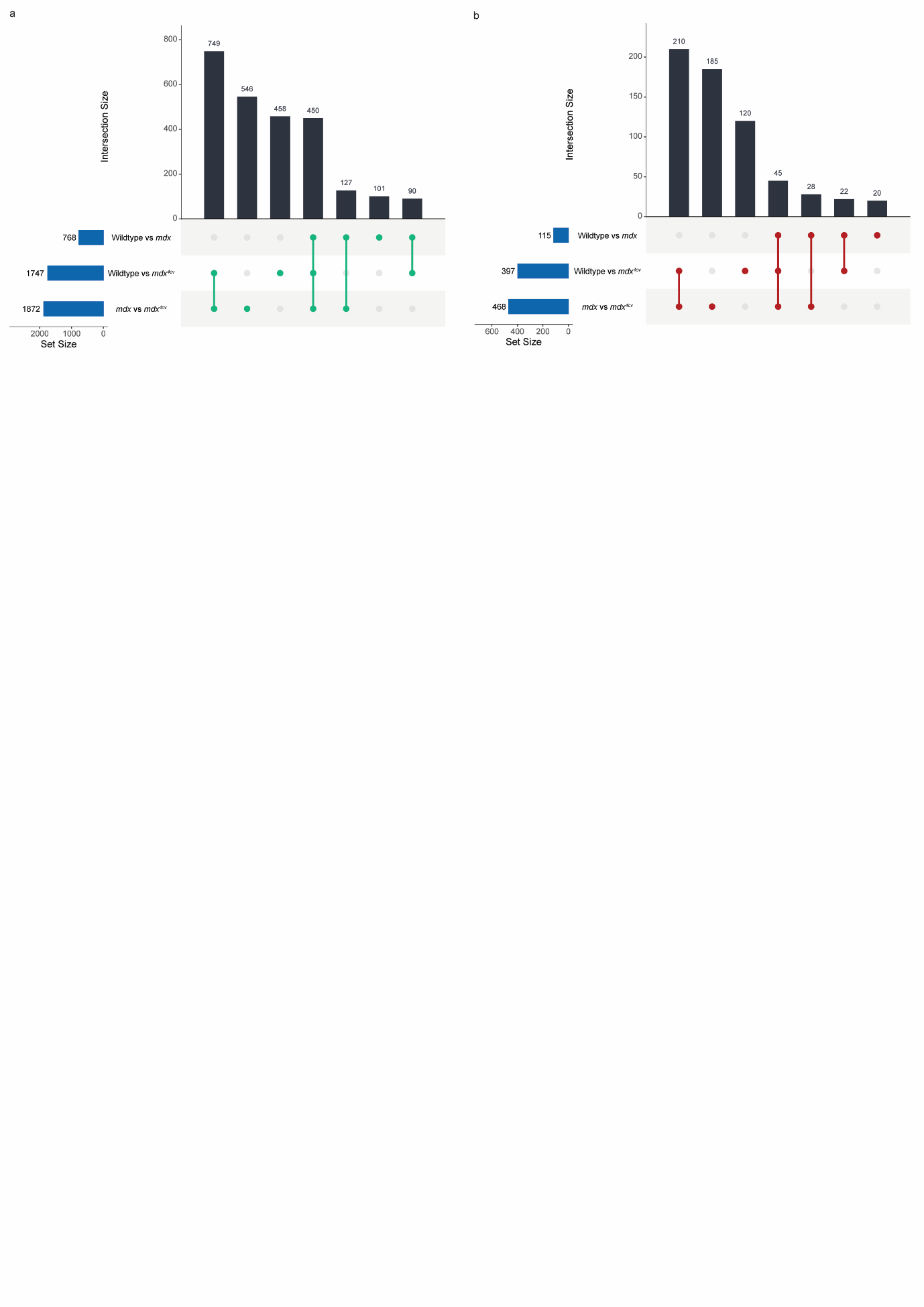

**Supplementary Figure 2. Intersection of differentially expressed genes (DEGs) across DMD mouse models.**

(a) UpSet plots illustrating the overlap of All DEGs between genotype contrasts across spatial brain transcriptomic data. encompassing any gene that reached significance (FDR < 0.05, |log2FC| > 0.1) in at least one brain region.

(b) UpSet plots illustrating the overlap of Core DEGs between genotype contrasts across spatial brain transcriptomic data. Core DEGs are defined by consistent significant differential expression in at least three distinct brain regions.

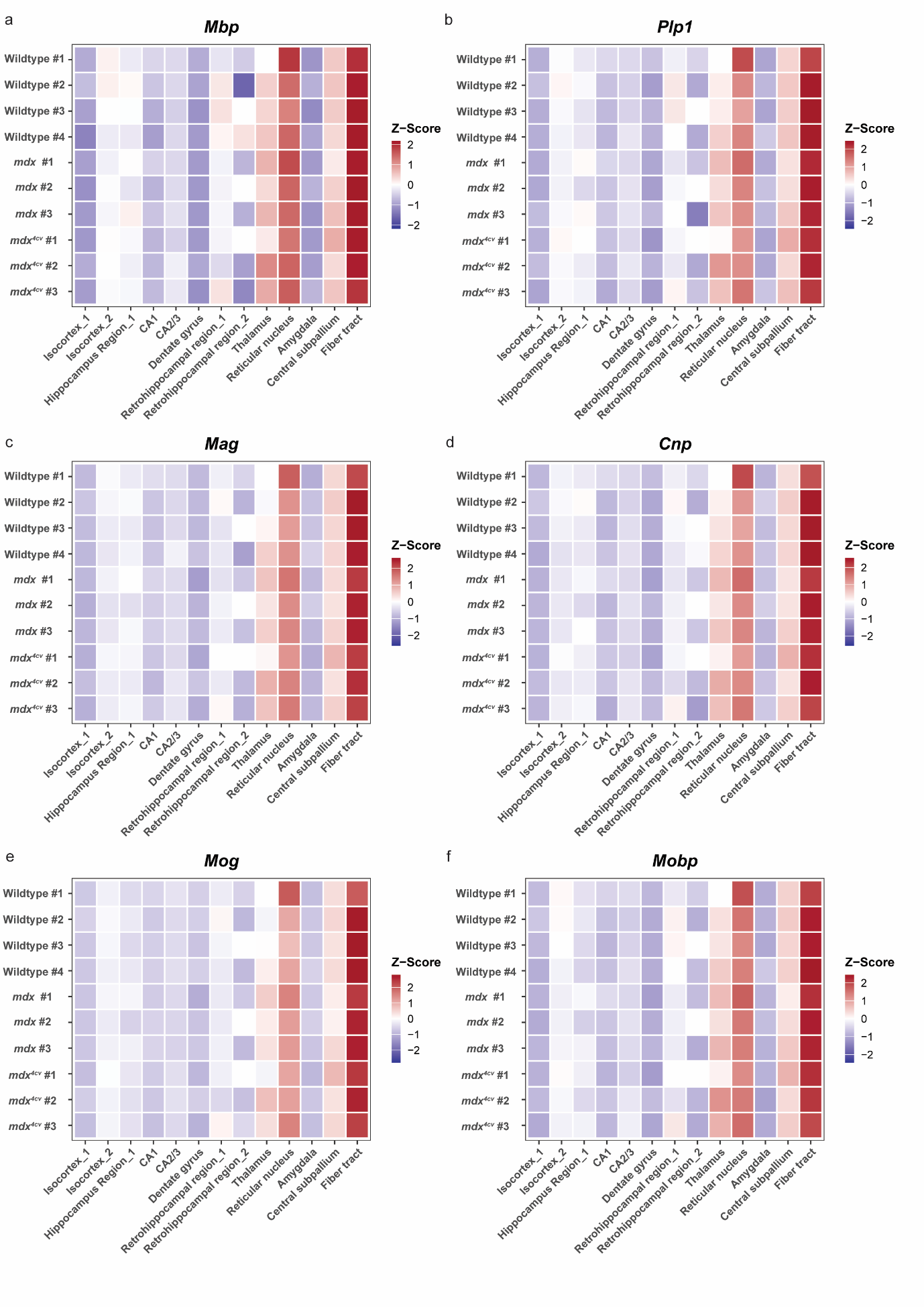

**Supplementary Figure 3. Regional expression patterns of myelin associated genes across wildtype, *mdx*, and *mdx^4cv^* mouse brains**

Heatmaps represent the Z-scaled average expression of myelin-associated genes (a) *Mbp*, (b) *Plp1*, (c) *Mag*, (d) *Cnp*, (e) *Mog,* and (f) *Mobp* across brain regions in WT, *mdx*, and *mdx^4cv^* mouse models. Expression values were averaged by region and scaled within each individual sample to highlight relative spatial enrichment (red) or depletion (blue) across region.

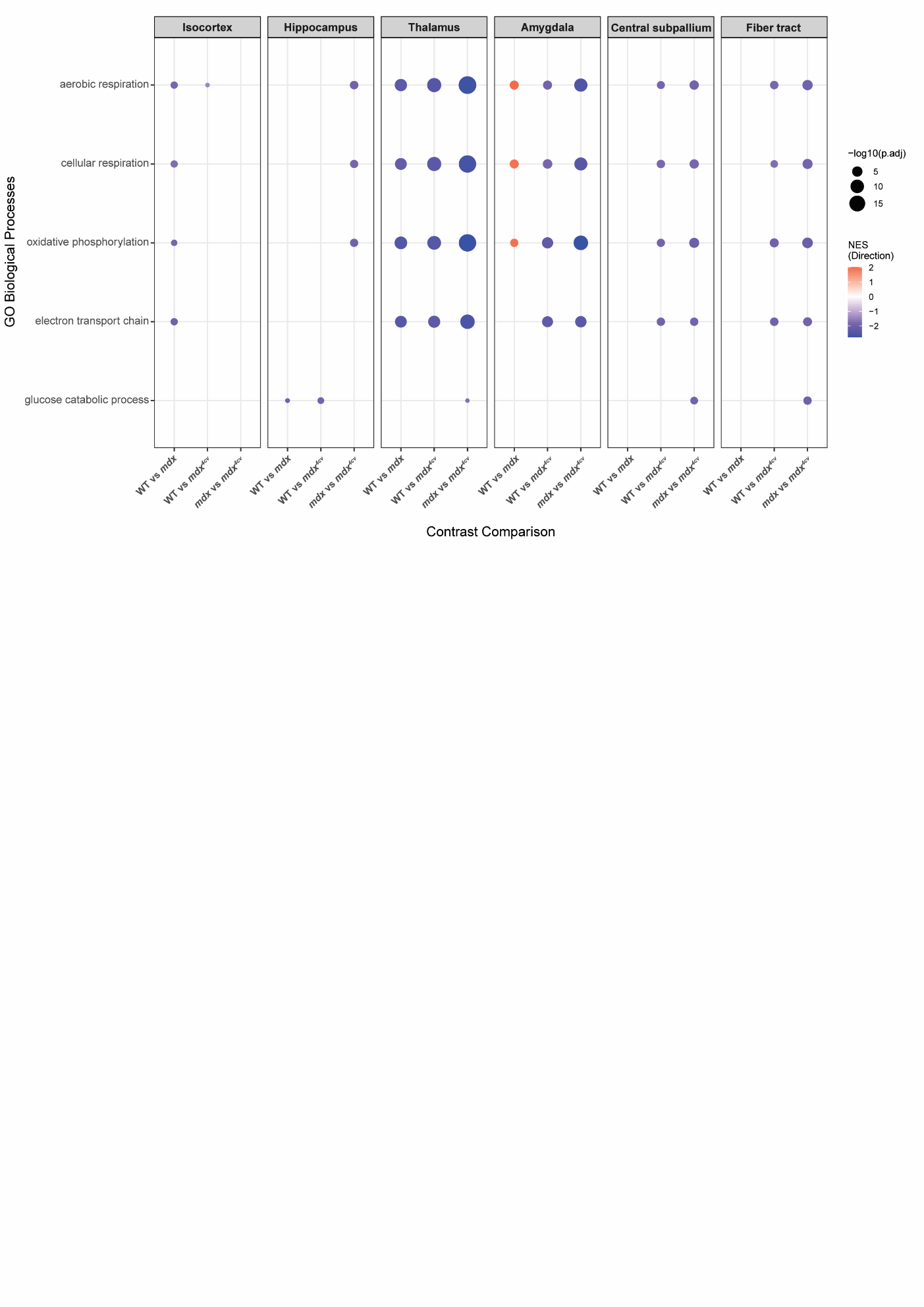

**Supplementary Figure 4. Regional enrichment of energy metabolism pathways.** Dot plot showing targeted Gene Ontology (GO) biological processes across distinct brain regions in wildtype, *mdx*, and *mdx^4cv^* mouse models. Dot size reflects the statistical significance of the enrichment (–log10 adjusted *p*-value, cutoff < 0.05). Dot color indicates the Normalized Enrichment Score (NES), where red denotes an overall increase in pathway expression and blue denotes a decrease.

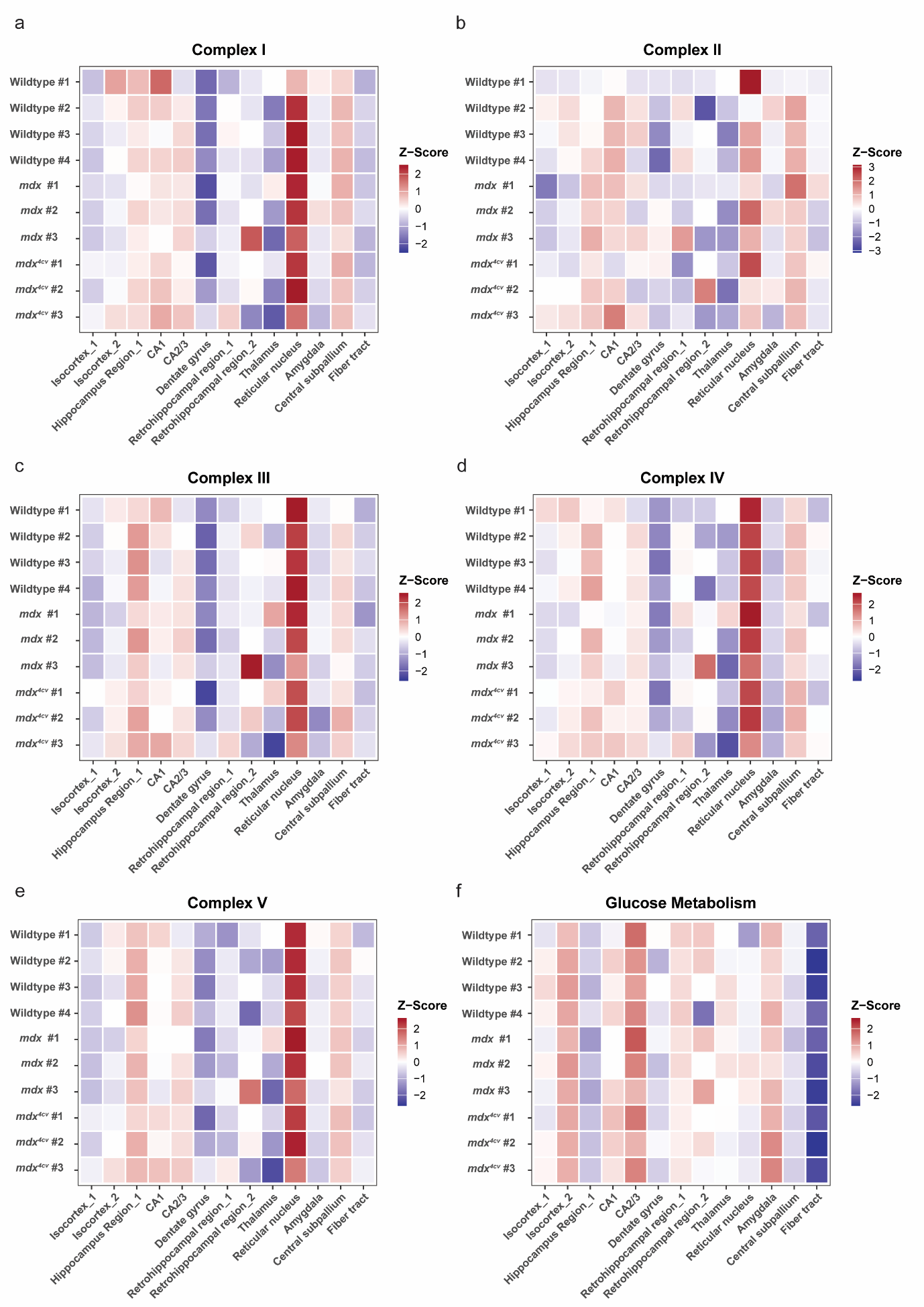

**Supplementary Figure 5. Regional alterations in each electron transport chain (ETC) complex and glucose metabolism pathways across wildtype, *mdx*, and *mdx^4cv^* mouse brains**

Heatmaps depict pathway-level expression signatures for ETC complexes (I-V) and glucose metabolism across brain regions in WT, *mdx*, and *mdx^4cv^* mouse models. Expression values were averaged by region and scaled within each individual sample to highlight relative spatial enrichment (red) or depletion (blue) across region.
